# A Generative Approach toward Precision Antimicrobial Peptide Design

**DOI:** 10.1101/2020.10.02.324087

**Authors:** Jonathon B. Ferrell, Jacob M. Remington, Colin M. Van Oort, Mona Sharafi, Reem Aboushousha, Yvonne Janssen-Heininger, Severin T. Schneebeli, Matthew J. Wargo, Safwan Wshah, Jianing Li

## Abstract

As the emergence of bacterial resistance is outpacing the development of new antibiotics, we must find cost-effective and innovative approaches to discover new antibacterial therapeutics. Antimicrobial peptides (AMPs) represent one promising solution to fill this void, since they generally undergo faster development, display rapid onsets of killing, and most importantly, show lower risks of induced resistance. Despite prior success in AMP design with physics- and/or knowledge-based approaches, an efficient approach to precisely design peptides with high activity and selectivity is still lacking. Toward this goal, we have invented a novel approach which utilizes a generative model to predict AMP-like sequences, followed by molecular modeling to rank the candidates. Thus, we can identify peptides with desirable sequences, structures, and potential specific interactions with bacterial membranes. For the proof of concept, we curated a dataset that comprises 500,000 non-AMP peptide sequences and nearly 8,000 labeled AMP sequences to train the generative model. For 12 generated peptides that are cationic and likely helical, we assessed the membrane binding propensity via extensive all-atom molecular simulations. The top six peptides were promoted for synthesis, chemical characterizations, and antibacterial assays, showing various inhibition to bacterial growth. Three peptides were validated with broad-spectrum antibacterial activity. In aggregate, the combination of AMP generator and sophisticated molecular modeling affords enhanced speed and accuracy in AMP design. Our approach and results demonstrate the viability of a generative approach to develop novel AMPs and to help contain the rise of antibiotic resistant microbes.

---

Antibiotic resistance, which arises when bacteria develop the ability to defeat small-molecule drugs designed to kill them, has become a severe threat to our society and industries. According to CDC’s Antibiotic Resistance Threats Report in 2019,^1^ over 2.8 million antibiotic-resistant infections occur in the US each year, over 35,000 deaths as a result. In fact, bacteria never stop finding ways to resist new antibiotics, and can even share their resistance with one another. Thus, persistent efforts are required to discover new molecules and to contain the spread of dangerous resistant bacteria. With advantages over traditional small-molecule antibiotics (e.g., broad-spectrum activity, rapid onset of killing, low levels of induced resistance, etc.), antimicrobial peptides (AMPs) show promise for treating infectious diseases caused by bacteria, viruses, or fungi,^2–5^ which are detrimental to our society and the healthcare, veterinary, and agriculture industries. Unlike traditional small-molecule antibiotics, AMPs are biological polymers predominantly containing 4 to 40 natural amino acids. So far, over 8000 AMPs^6^ have been found in animals and plants as a first-line defense of the host immune systems. However, the known AMPs represent only a small fraction of the vast chemical space — e.g., 20^*N*^ possible chemical compositions for a *N*-residue peptide composed of 20 natural amino acids. Recent methods have focused on quantitative structure activity relationship (QSAR) and machine learning (ML) predictions in order to seek new AMPs (see recent reviews^7–8^). Such approaches often rely on sequences to be generated randomly or evolved from known sequences,^9–14^ which can be computationally demanding to afford reasonable sampling in the vast peptide sequence space. Instead, it is more efficient to utilize a generative model to create new peptide sequences from the underlying distribution of possible AMPs. However, a prior study based on a variational autoencoder (VAE) model^15^ showed only two confirmed AMPs out of 20 predicted peptides, suggesting that improvements remain needed in the design accuracy to lower false positive rate and higher activity of generated AMPs. In this work, we have developed, for the first time, a generative adversarial network (GAN) conjugated with molecular modeling to design AMP-like sequences, which is fundamentally distinct from previous AMP design approaches. Validated by experiments, we have obtained three AMPs out of six tested peptides, which indicates the potential of our approach to be applied as a useful tool for future AMP discovery.

Our AMP-GAN approach (Figure 1) utilized a modified conditional generative adversarial network (CGAN).^16–17^ CGAN represents a powerful approach for creating generative models, as it gives instructions to two networks that are pitted against each other in a zero-sum game. We have utilized this framework and a set of only four fundamental conditions (or labels) to generate new AMP sequences, in contrast to the large number of descriptors in prior QSAR studies.^10, 13^ The discriminator was trained to distinguish between authentic and generated sequences, and the generator was tasked with generating sequences that fool the discriminator. In order to trick the discriminator successfully, the generator must learn the underlying AMP distribution. By generating sequences from the underlying AMP distribution and providing conditioning variables that allow the generation process to be directed, we are able to exert control over the properties of generated sequences. Furthermore, the conditional vectors provided a distinct capability to target particular pathogens, mechanisms, and even sequence lengths. This directed generation approach moves machine learned AMP discovery away from a rejection sampling methodology and towards precision design, which allows for increased accuracy towards discovering novel AMPs with desirable activity, selectivity, and safety profile.

**Figure 1.**
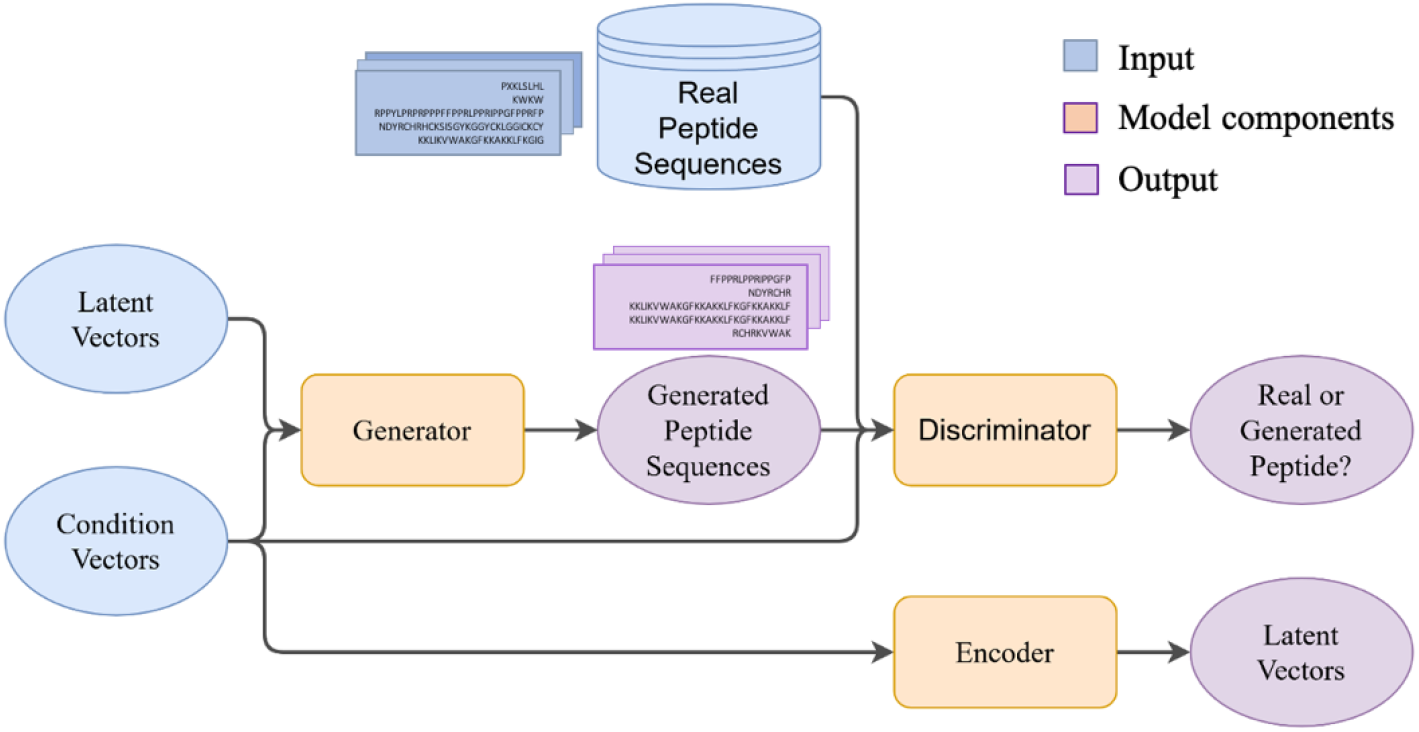
Illustration of the organization of AMP-GAN.

With a large number of AMP-like peptides designed by a generator, it is essential to assess their structures and potential interactions, for example using molecular modeling and simulation. Recent advances in force fields^18–19^ and solvent models^20–21^ have greatly enhanced the accuracy of peptide modeling. For example, current tertiary structure prediction can obtain near-native models for up to 95% of peptides within 52 residues in length.^22–24^ Also, molecular dynamics (MD) simulations have provided valuable structural and mechanistic insight into peptide interactions with biomolecules, including AMP aggregation in solution and membranes^25–27^, AMP-induced membrane disruption and poration,^28–39^ etc. In particular, free-energy simulation techniques have become available for reliable semi-quantitative or quantitative assessment for membrane binding,^40^ membrane permeation,^41^ and membrane pore size,^42^ as well as peptide mutations.^43^ Therefore, we carried out free-energy simulations to verify the potential interactions of the AMP candidates with a lipid bilayer membrane, and estimated the relative membrane binding propensity. Our experimental validation indicates the success of our *in silico* approach to combine sequence-based generation and structure-based ranking/selection. In aggregate, with a comprehensive consideration of both AMP sequences, structures, and interactions, our approach and discovery may open a novel avenue for future AMP discovery.

## Results

### 1. AMP-GAN generated peptides with high sequence diversity

We constructed AMP-GAN to produce a large number of AMP-like sequences. AMP-GAN was trained over a data set comprised initially of 8,005 known AMPs from the DBAASP^6^ and AVPdb^44^ databases as well as nearly 500,000 non-AMP peptides selected from the Uniprot database.^45^ Four conditions were chosen based on their fundamental nature and data abundancy: sequence length, microbial target, target mechanism, and MIC50 (the concentration of peptide which inhibit 50% of bacterial growth). Distinct from prior research, we designed two novel labels regarding the microbial target (Gram-positive/negative bacteria, fungi, viruses, or others) and the target mechanism (e.g., disrupting the lipid membrane, inhibiting vital proteins, or interfering with DNA/RNA), which allowed us not only to train our network on the broadest set of AMPs, but also to capture the potential connections between them. As a result, each generated sequence was associated with specific microbial target(s) and potential mechanism(s), which greatly facilitated subsequent selection and experimental validation. As a matter of practice, we focused on AMPs within 32 amino acids in length due to the affordable synthetic cost. Thus, we applied a maximal 32-residue length constraint to both the training sequences and generated sequences (Figure 2A). Notably, while helicity or secondary structures were commonly included in the descriptors in several QSAR models,^13–14, 46–47^ there is no structural constraint in our AMP-GAN. Thus, AMP-GAN can produce novel peptide sequences that fold into different three-dimensional (3D) structures.

**Figure 2.**
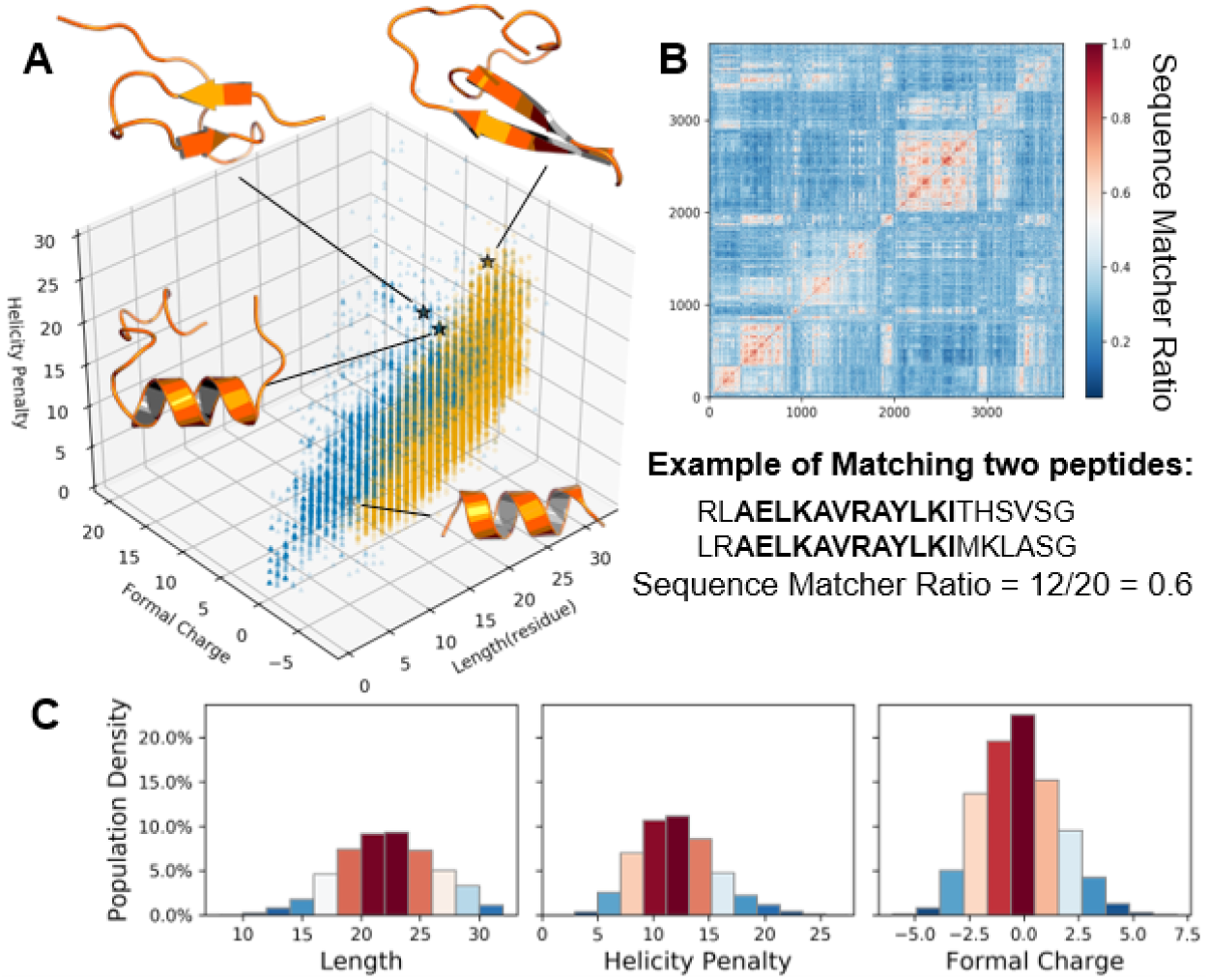
Diversity of AMP-GAN-generated peptide sequences and structures. **(A)** Comparison of the training set (blue dots) and the generated data set (orange dots) in terms of peptide length (in residue), formal charge (in electron charge unit), and helicity penalty (in kcal/mol). **(B)** Heatmap to show the sequence diversity among the generated peptides of 20 residues in length (3829 peptides in total). X-Y axes represent numbering of the 20-residue peptides in a list. The long contiguous matching subsequences were identified between two given peptides (see the example). A ratio of 1 shows exact identical sequences (the diagonal in the heatmap), while the sequence matcher ratio remains low among most of our generated data, an indicator of sequence diversity within our data set. **(C)** Histograms to show the distribution of peptide length (in residue), formal charge (in electron charge unit), and helicity penalty (in kcal/mol) in our generated data set.

AMPs are diverse in sequences and structure, and therefore an important indicator for a good AMP generative model is the reproduction of generated sequences similar to known AMPs, but unique enough to be novel. Thus, it is a major challenge for many models trained on relatively scarce and sparsely labeled data to generate new sequences that are distinct, both from known sequences and from each other in the data set. In this work, we generated 50,000 sequences with AMP-GAN using condition vectors drawn from the training data (known AMPs and known non-AMPs). We compared the training data (known AMPs) and generated data (AMP candidates) in terms of length, formal charge (FC), and helicity penalty (HP) as shown in Figure 2A. HP was determined by the sum of energy scores^48^ associated with the helix propensity of each residue, e.g. no penalty for alanine as (0 kcal/mol) and high penalty for glycine (1 kcal/mol) and proline (3.6 kcal/mol). Overall, the 50,000 AMP candidates displayed normal distributions in terms of length, FC and HP (Figure 2C.). As shown in Figure 2A, the two distributions overlap sufficiently to imply a consistency between generated peptides and known AMPs. To assess the sequence diversity within the AMP candidates, we calculated the pairwise similarity using the ratio from *SequenceMatcher* in the difflib python package (https://docs.python.org/3/library/difflib.html), which was defined as the fraction of contiguous matching subsequences between two given peptides. With the 3829 20-residue peptides as an example, the majority of peptides are distinct from each other, as indicated by the low sequence matcher ratio (Figure 2B). In addition, low similarity was found between our generated sequences and known proteins and peptides in the UniprotKB^45^ database, using the Basic Local Alignment Search Tool (BLAST). For sequences below 10 residues in length (Table S1), we found 73% of our AMP candidates with accidental homology to regions of larger proteins (E-value 0.1-10), while the rest were novel sequences unrelated to known proteins (E-value >> 10). For longer AMP candidates (10 to 32 residues), we estimated that no more than 3% of sequences generated by AMP-GAN had significant homology (E-value < 0.1) to known peptides or proteins. Generally, these analyses indicate a strong ability of AMP-GAN to generate a diversity of novel AMP-like sequences and to explore new areas in the vast peptide chemical space.

### 2. AMP-GAN-generated peptides with high structural diversity

Generation of diverse AMP structures is notably more difficult than the generation of sequences alone. However, the sequence diversity from AMP-GAN does serve as a firm foundation for the generation of structurally diverse AMP candidates. Although the vast structural diversity of AMPs has been long known, currently the ~1200 known structures of AMPs (collected in APD3 ^49^ until June 2020) show that near 1/3 of the AMP structures are classified as helical. Most helical AMPs are believed to exert bactericidal effects via irreversible rupture of bacterial membranes, while a small but increasing number of non-helical AMPs are revealed to modulate vital bacterial proteins.^50–52^ This suggests a critical need to explore new areas in the chemical space and to seek not only helical but also non-helical AMP designs. Most prior studies of AMP design were restricted to α-helices or simply ignored the peptide structures.^10, 13, 53–54^ Distinct from these studies, the generator of our AMP-GAN, while not explicitly trained on information about secondary or tertiary structure, can infer through implicit sequence-structure relationships the underlying diversity of AMP structures. Thus, AMP-GAN is allowed to generate a broad range of peptide sequences, which likely fold into diverse 3D structures as well as result in various potential interactions and selectivity against the pathogenic targets.

Using peptide structure prediction,^55^ we showed that the generated sequences likely adopt various folded structures such as *α*-helices, *β*-hairpins, and other prototypical motifs (Figure 3A). Focusing on peptides shorter than 32 residues, the structural diversity observed in selected cases was in agreement with what was expected from known AMP structures. For example, several sequences with high HP (HP/res > 0.7) were more likely to adopt stable hairpin structures, which were confirmed by 10-ns MD simulations. In contrast, almost all the peptides with low HP (HP/res < 0.5) formed helical structures. Thus, HP provided a simple, fast estimation of the AMP secondary structures in our studies (Figure 3B). Notably, natural hairpin AMPs typically contain one to three disulfide linkages.^56^ Although we did not include the redox information in the current design of AMP-GAN, our MD simulations confirmed that several AMP candidates could adopt stable b-hairpin motifs, even in the absence of disulfide bonds (Figure 3A). With the high structural diversity of the generated peptides, AMP-GAN can be a useful tool to identify new AMP-like sequences and to allow ad hoc studies to model the structural stability, design chemical modifications, and predict potential interactions with key components (like membranes, vital proteins, and DNA/RNA) of the pathogen cells.

**Figure 3.**
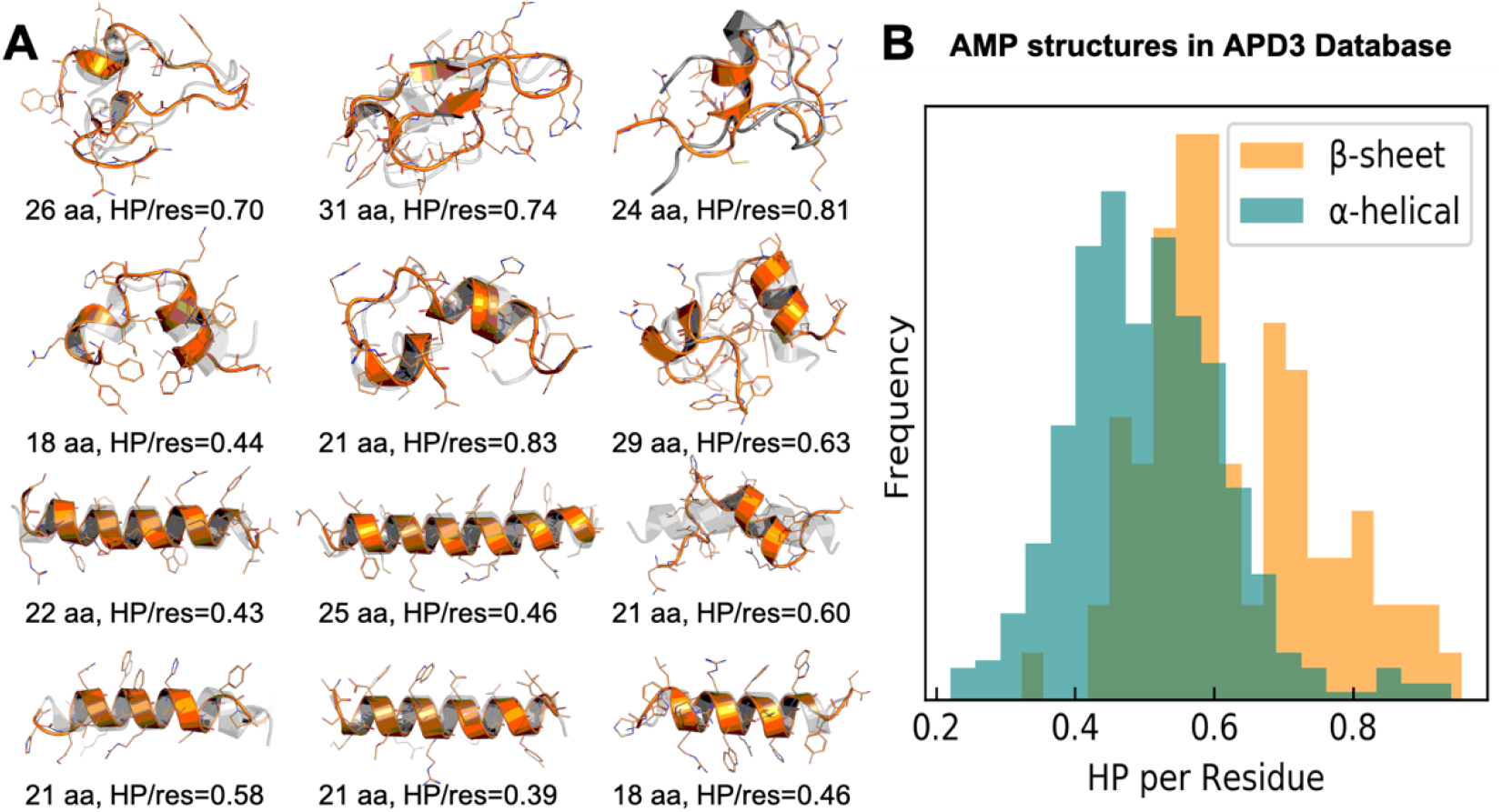
**(A)** Predicted structures of 12 cationic AMP candidates (formal charge > +3). Each structure in the orange cartoon is from the final snapshot of a 10-ns MD simulation. The structure in grey is the initial model that is either predicted by Pep-Fold55 or restricted to be helical. The peptides with low HP/res are more likely to adopt stable helical structures while the peptides with high HP/res may fold into *β*-hairpin structures (without disulfide bonds) or other motifs. **(B)** Distribution of known AMPs that are classified as α or β structures in terms of HP per residue, which included ~400 *α*-helical and ~80 *β*-hairpin AMPs.

### 3. Molecular modeling to select potent helical AMPs against bacteria

We first obtained 50,000 AMP-like peptides from AMP-GAN (Figure 1A), from which we selected a smaller set of candidates for synthesis and experimental validation. Compared with experiments, molecular modeling and simulations are still affordable with a relatively large number of AMP candidates and thus can be useful to eliminate false positives from AMP-GAN and improve our accuracy in the selection process. To show the effectiveness of molecular modeling, we focused on *α*-helical AMP candidates potentially against Gram-positive/negative bacteria, the main mechanism of which has been well accepted as membrane disruption and interaction.^57^ Based on the information provided by AMP-GAN (activity, sequence charge, HP, etc.), we narrowed down to 12 AMP candidates (sequences and conditional labels shown in Table S2), six of which were further identified by a free-energy simulation approach to estimate the membrane binding propensity, which is described as follows.

Firstly, we eliminated all sequences which had targets against mammalian, cancer, or fungal groups in their conditioning vector. We further excluded all peptides that had labels above the lowest band of predicted MIC50 activity (~5.76 μg/ml) from our conditioning vectors, in order to keep the most potent AMP candidates. Then, two selection criteria were applied to identify chemically relevant candidates for antibacterial tests. (i) *The structure rules*: a total formal charge greater than +2 or a calculated HP < 5 kcal/mol;^48^ (ii) *The chemical rules*: elimination of sequences with multiple adjacent Ser or Gln or ones which had selenocysteine or pyrrolysine and sequences which did not contain adjacent Arg and Trp (known as the RW pattern commonly seen in many AMPs). The structural rules were used to select sequences likely to be charged or helical which would be easier for us to confidently model the peptide structures and interactions, while the chemical rules helped identify chemically relevant antibacterial peptides that avoided unstable and difficult to synthesize peptides. At the end, 12 sequences passed all of these filters.

To quantitatively rank the 12 candidates and select the most-likely membrane-active peptides, we carried out all-atom free-energy simulations and calculated the membrane-binding propensity. For each peptide, an estimate of the free energy change Δ*G* (from embedding in a model *E. coli* bacterial inner membrane) was obtained via a quick set of MD simulations with umbrella sampling (US).^58^ US restrained the helical peptides to different heights above and below the membrane surface, and used the weighted histogram analysis method to estimate Δ*G*. The determined Δ*G* for each peptide was used to obtain a score correlated to the membrane-binding propensity, given by Equation 1, which compares a candidate sequence to a known membrane-active peptide, magainin 2 (Δ*G*_+_) and a known non-AMP sequence (Δ*G*_−_, sequence in the SI).

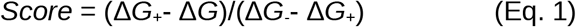

In Equation 1, a positive (or negative) score represents an AMP sequence with a higher (or lower) propensity to bind the membrane than magainin 2. This difference is then scaled by the range of the positive and negative controls for relative comparison. US was repeated in three replicas for the control sequences and showed they have significantly different Δ*G* values with Δ*G*_+_ 7 kcal/mol lower than Δ*G*_−_ and a 5-13% relative error (calculated ΔG values in Table S3), suggesting this methodology can distinguish poor from good membrane binding peptides. It is noted that the significance of our free-energy calculation is in the relative values, which is confirmed by the control sequences following in the expected trend that a known AMP would have a more negative change in energy upon membrane binding than a non-AMP (i.e. known AMPs have a more favorable change in energy upon binding to anionic membranes). Based on their membrane-binding propensity scores, we ranked the 12 helical peptides (Table S3) and decided to promote the top six peptides for synthesis and experimental validation (Table 1).

**Table 1.**
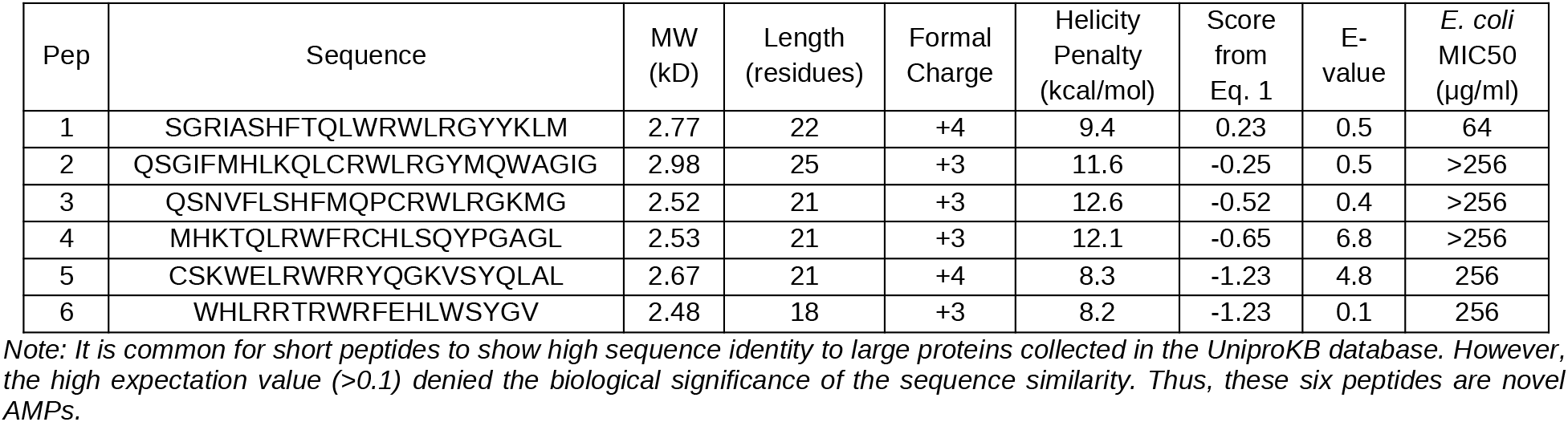
Properties of six selected AMP candidates. The E-value and sequence identity were obtained from BLAST, while MIC50 values were measured by serial dilution growth inhibition assays. All the peptides are amidated in the synthesis.

### 4. Experiments to validate activity and toxicity of AMP candidates

The six peptides with the most preferred membrane-binding propensity were purchased from Genscript (> 90% HPLC purity) and tested against four microbes, Gram-negative *E. coli*, *K. pneumoniae*., *P. aeruginosa*, and Gram-positive *S. aureus*, in order to validate our methodology. We used a serial dilution method from 256 μg/mL to 0.01 μg/mL in an attempt to find the value for which approximately 50% of the bacterial growth was inhibited (MIC50). Almost all of the six peptides show growth reduction to different extents against Gram-positive and -negative microbes (Figure 4). Their antibacterial activity increased with the increasing concentration, but increases in light scattering (due to peptide aggregation) might also increase the apparent bacterial growth at high concentrations. While this was partially controlled for by subtracting peptide-alone optical density from corresponding growth measurements, this effect does reduce the accuracy of the assays at higher peptide concentrations. For *E. coli*, Pep1 at 64 mg/ml and Pep5 and Pep6 at 256 *μ*g/ml inhibited over 50% of bacterial growth, while the others achieved less than 40% inhibition at the concentration below 256 *μ*g/ml. For *S. aureus*, four of the six peptides inhibited ~50% bacterial growth at the concentration of 256 *μ*g/ml, except Pep3 and Pep4. In addition, most of our peptides inhibited *P. aeruginosa* and *K. pneumoniae* growth by 20-30% at a concentration below 256 *μ*g/ml (Figure S4). Interestingly, AMP specificity was achieved with Pep3 which targeted *S. aureus* but not the Gram-negative bacteria which we tested (Figure S4), while more broad-spectrum activity was observed for other peptides. In general, the AMP-GAN model produced novel and active peptides against the variety of bacteria tested. Two of the most interesting peptides, Pep1 and Pep6, were further tested for toxicity with A549 human cells. At concentrations below 300 μg/mL, both Pep1 and Pep6 had near 90% viability compared to our control. At 300 μg/mL, Pep1 displayed notable toxicity with less than 5% of the cells still viable compared to our control, but the less potent Pep6 appeared to be less toxic with near 90% viability (Table S4).

**Figure 4.**
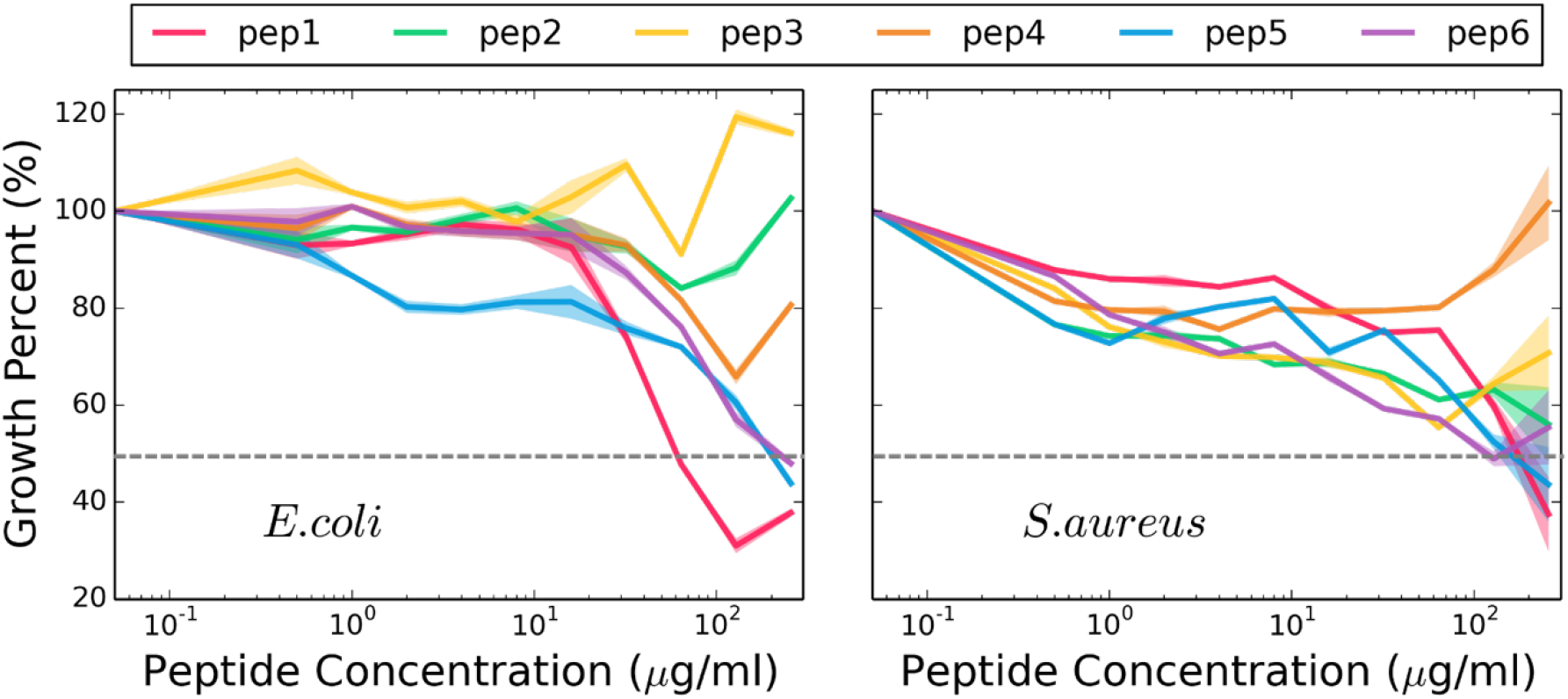
Bacterial growth assays for the six synthesized peptides against *E. coli* (left) and *S. aureus* (Right). The growth of the bacteria was measured by optical density at 600 nm (OD_600_) and normalized as a percentage to bacterial growth with no peptide. These normalized values are plotted against AMP concentration. Dashed lines to judge MIC50 concentrations are shown. Error bars are from three independent experiments. Each experiment was carried out with 3 replicas, with shaded error bars.

Our experimental MIC50 are higher than the predictions from AMP-GAN, which may be impacted by several chemical and biological factors. For example, Pep2 and Pep4 with low solubility (< 5 mg/ml in water) likely aggregate in the aqueous solution, which reduces the effective peptide concentrations. Further, despite the helical assumption, some of these peptides may undergo significant conformational changes during the membrane binding process, which can affect our selection accuracy. Similarly, the AMP-GAN generated sequences of length that varied from the input label (R^2^=−1.37). Length is the only label that can be verified on a large-scale without experimentation. This poor correlation implies that while the AMP-GAN in this current version may have learned the underlying distribution, it still needs to better interpret the significance of the labels — a direction which we will pursue to improve AMP-GAN. In addition, it has been reported that some Gram-negative bacteria can secrete outer membrane vesicles to prohibit AMP acting on the cellular membrane.^59^ Despite various impacts to the actual AMP activity, our best peptide, Pep1 has comparable broad-spectrum bactericidal activity to some natural AMPs like amphibian AMP magainin 2 and human AMP LL-37, which demonstrates the potential of our approach to rationally design active AMPs.

## Discussion

### 1. Synergy of the sequence-based generation and structure-based modeling

There are two distinct ways to achieve AMP sampling with machine learning — ***classifiers*** (which provide a yes/no answer for a given peptide sequence) and ***generators*** (which directly provide new predicted active sequences). While classifier models^9–10, 13–14, 54^ are often more accurate than generator models,^15, 58^ they can be inefficient to discover diverse AMP candidates and requires lots of human expertise and effort to construct the descriptors when combined with QSAR. Compared with the classifiers, AMP-GAN can be more efficient to design AMP-like sequences according to an approximation of the underlying distribution in the AMP chemical space. Our review of current classifier and generator approaches suggested a trade-off between sampling diversity and prediction accuracy: restricted AMP sampling can achieve higher accuracy to identify active AMP-like sequence, while AMP-GAN and another deep generative model (VAE) can produce much more novel sequences, with the prediction accuracy to be improved.

In this work, our combination of sequence-based generation (AMP-GAN) and structure-based modeling (US sampling) offers a solution to the trade-off. First of all, a feature in AMP-GAN, is that useful information (like potential activity, potential pathogenic target, etc.) is included in the design of peptides for specific tasks (antibacterial, antifungal, anticancer, etc.). The subsequent molecular modeling (especially free energy simulations), as shown in this study, is useful to quantitatively rank the AMP candidates. Compared with a prior study using conventional MD simulations,^15^ our free-energy simulation approach is more effective to reduce the high false positive rate. Our overall approach is fundamentally distinct from previous ones which employed both sequence- and structure-dependent descriptors or labeling. In principle, the limited number of labels allow for a greater focus on purely exploring the sequence space as labels do not need to be fine-tuned. Furthermore, we can obtain a higher accuracy in selection of AMP candidates with simulations based on molecular structures and interactions. Last but not least, the sequence-based generator and structure-based scoring are well integrated. Although in this work, we provide the proof of principle to seek helical AMP to target the membranes of bacterial pathogens, our approach, with some fine tuning of the AMP-GAN in order to better observe labels and to ensure more stable training, is ready to be used to study helical or non-helical AMPs with many other therapeutic targets (like proteins and DNA/RNA) beyond bacteria.

### 2. Potential molecular mechanism of Pep1

The 22-residue Pep1 was shown to be the most active AMP in our design, and its sequence is novel with only accidental similarity to larger proteins (fragment < 10 residues). We further examined the interactions between Pep1 and lipids in our simulations and via Circular Dichroism (CD) spectra (Figure 5). It is shown by CD that without the surfactant sodium dodecyl sulfate (SDS), Pep1 was only 7% helical and largely unordered in solution. With the presence of SDS, Pep1 dramatically increased its helicity to 74% in the SDS micelles, which was consistent with our observation from MD simulations (55-77% helicity). Analysis of our simulations provided the molecular detail of Pep1 interactions with a lipid bilayer model.

**Figure 5.**
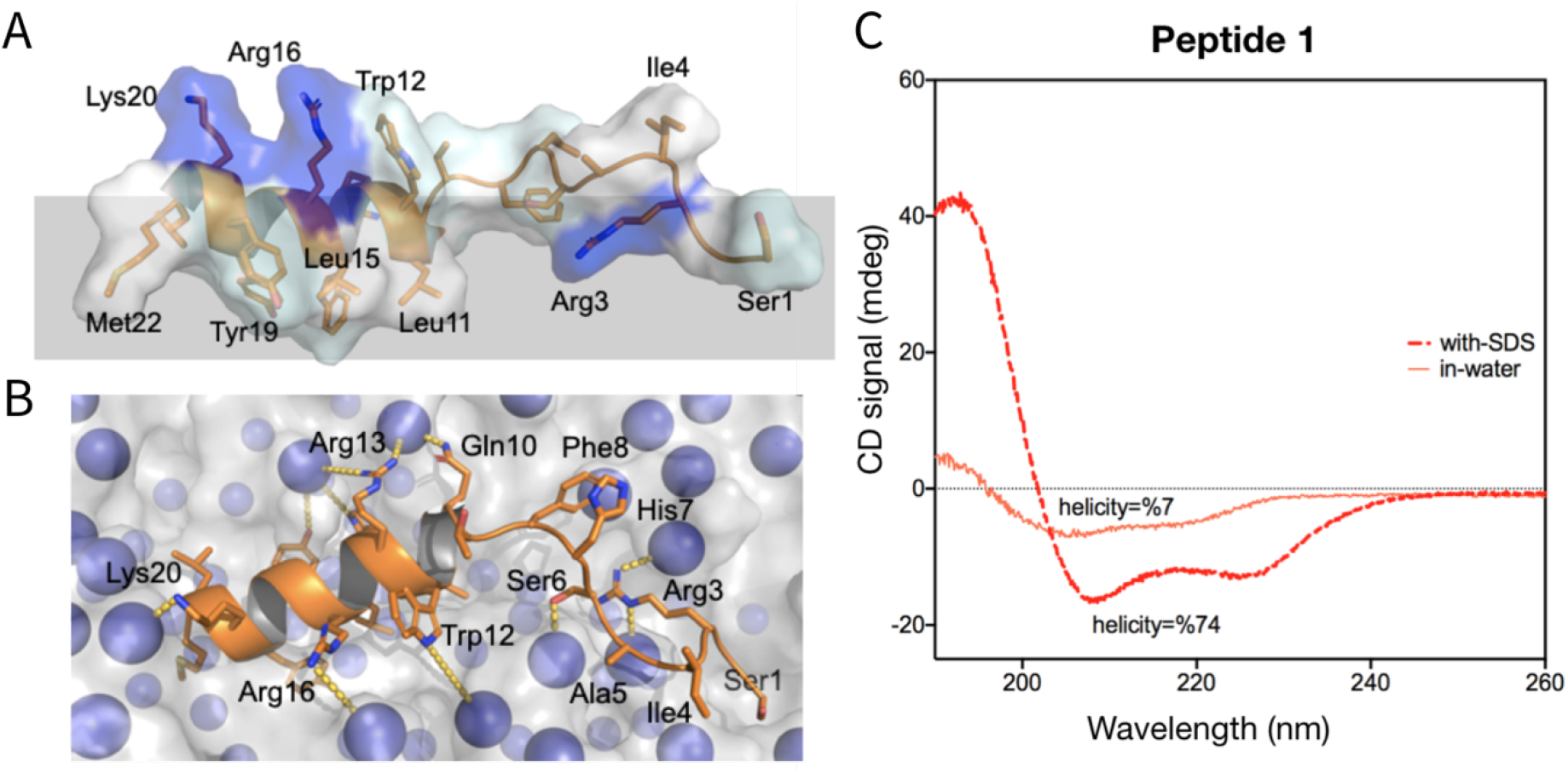
Membrane binding motifs and solution structures of Pep1 **(A)** side view, **(B)** top view. **(C)** CD spectra of Pep1 with and without SDS.

The helical conformation of Pep1 resembles a typical amphipathic helix, with a relatively hydrophobic face (Leu11, Leu14, Tyr19, and Met22) and a cationic face (Arg3, His7, Gln10, Arg13, Arg16, and Lys20). In our simulations, the hydrophobic sidechains on helical Pep1 were embedded in the non-polar environment of the membrane core (Figure 5A-B) This orientation coincided with charged sidechains interacting with the charged phosphate groups on the membrane surface, establishing a combination of strong interactions on all sides of the helical segment. Taken together, Pep1 likely undergoes a disorder-to-helix transition during membrane binding. The folding of Pep1 results in an amphipathic helix, with strong interactions with the anionic lipids in the membrane. Supported by collaborated simulation and spectroscopic evidence, Pep1 likely adopts a mechanism of a typical helical AMP like melittin. Interestingly, we did not observe the other five peptides which we synthesized with a similar transition like Pep1 (Figure S5). However, the various level of helicity of these peptides indicates the capacity of AMP-GAN to generate structurally diverse AMP-like sequences.

## Concluding Remarks

In summary, we have invented and demonstrated an efficient and accurate AMP design methodology. By curating a dataset that comprises ~500,000 non-AMP peptide sequences and ~8000 labeled AMP sequences to train the AMP-GAN model for generating new AMP sequences, we created a general generator that can be used to target more than just one type of microbe or one mechanism. We then demonstrated a proof-of-concept method for evaluating peptide candidates by evaluating membrane binding tendency toward a model *E. coli* inner membrane. This technique can be extended to different membrane compositions and therefore different microbes, thus retaining the generality of our generator in our screening as well. This synergy of sequence-based generation and structure-based ranking represents a new methodology toward flexible, precision AMP design. Extensive analysis of the generated antimicrobial sequences reveals that the proposed framework, beyond being general, is indeed capable of learning and generating from a richer representation than trained upon and yields AMPs that are both diverse in sequence and structure. Our methodology as an entity will be valuable in the ever-necessary rational design of AMPs with the potential to be expanded to even broader classes of biologics.

## Materials and Methods

### 1. Computational Methods

We utilized a GAN approach that incorporates several elements from recent research. The base of our model is a Wasserstein GAN with Gradient Penalty (WGAN-GP), which claims to mitigate many of the pathological learning dynamics that can occur when training GANs, including dissociation and mode collapse. Next, we add conditioning information following the structure outlined by Conditional GANs (CGAN), allowing human designers to influence the features of the generated peptide sequences. Additionally, we include an encoder component following work on Bidirectional GANs (BiGAN), which greatly simplifies the implementation of latent space interpolations and allows for more effective exploration of the learned latent space through the use of known AMPs. Figure 5 provides a flow diagram that depicts the high-level organization of our AMP-GAN, which is similar to what was investigated by Perarnau *et. al.* ^60^, excluding the label/conditioning vector encoder. Architecture details for the generator, discriminator, and encoder networks are shown in Figs. S1, S2, and S3 respectively.

For our training data we utilized the Database of Antimicrobial Activity and Structure of Peptides (DBAASP) as our known good AMPs ^6^. For those AMPs with multiple MIC50 readings we converted all readings into μg/mL and averaged them these were then binned into 10 deciles and those deciles which were provided to the AMP-GAN. We also combined categories of different microbes into a reduced set of Gram-positive, Gram-negative, Viral, Fungal, Mammalian, and Cancer. Similarly, we collapsed targets into, Lipid Bilayer, RNA/DNA, Cytoplasmic Protein, Membrane Protein, and Virus Entry. We also removed any sequences of length greater than 32 amino acids. For any sequences with a wildcard FASTA symbol (X, B, Z, or J) the symbol was replaced with a randomly selected concrete symbol during training. For our null data set we selected sequences from the Uniprot database, filtering any sequences that were present in DBAASP or were longer than 32 residues. The conditioning vectors for our null database were constructed to indicate no target microbes, no target mechanisms, maximal MIC50, and the appropriate sequence length. We trained AMP-GAN on 100,000 batches with 128 samples per batch. Each batch was composed of 64 randomly sampled AMPs and 64 randomly sampled non-AMP peptides. This translates to approximately 1,000 epochs of training over the positive dataset and 13 epochs over the negative dataset. Implementation details for the data processing, model creation, and model training elements can be found on [GitLab](https://gitlab.com/vail-uvm/amp-gan/-/tree/test_samples/).

For all atom molecular dynamics simulations, initial peptide structures were modeled as a helix and translated 25 Å above a 3:1 mass-ratio POPE:POPG membrane. The CHARMM-36m ^18^ forcefield was then applied using CHARMM-GUI ^61–62^ and the systems solvated with TIP3P water model and 120 mM NaCl. All simulations were performed using the AMBER18 simulation package ^63^. The following US protocol was developed to enable rapid screening and estimation of the relative free energy change upon peptide binding to the model membrane. Each peptide was put through the multi-stage minimization and equilibration protocol. Short, 500 ps, steered molecular dynamics (SMD) trajectories were run with a very stiff spring constant of 500 kcal/molÅ applied to a custom collective variable representing the z-component of the center of mass position of the peptide, over a total distance of 80 Å into the membrane. While this distance was larger than the membrane thickness, it was required to embed the peptide into the membrane over the short period due to translation of the membrane during SMD. Following SMD, frames most closely representing eight umbrella sampling windows linearly spaced between 20 angstroms below the phosphorous atoms in the top leaflet of the membrane and the initial position above the membrane were selected. To allow the membrane atoms and peptide sidechains to relax from the SMD simulations each window was equilibrated for 1 ns using a stiff spring constant of 1 kcal/mol Å. Following this period, 20 ns of production umbrella sampling simulations were performed with a weaker spring constant of 0.1 kcal/molÅ that ensured proper overlap of the umbrella windows. The final data was analyzed using the weighted histogram analysis method, and the final difference between the first and last window were used to estimate the free energy difference for embedding the peptide in the membrane.

### 2. Experimental Methods

These peptides were purchased from Genscript and characterized for solubility and helicity. Helicity was determined by circular dichroism (CD) spectra with a peptide concentration of 20μM in 1mL Milli-Q Water. The Jasco J-815 spectropolarimeter was set to analyze the peptides at a wavelength range of 260-190nm, a cell length of 2mm, and 3 scans. The resulting spectra was then fed to Dichromweb ^64^ and the helicity was then calculated using the CDSSTR method and a reference dataset.

Bacteria were stored as 15% glycerol stocks at −;80 °C and routinely propagated on LB agar or broth (Lysogeny broth, Lennox formulation). The specific species and strains used for these assays were *Escherichia coli* K12, *Pseudomonas aeruginosa* PAO1, *Klebsiella pneumoniae var. pneumoniae* KPPR1, and *Staphylococcus aureus var. aureus* ATCC 12600. For antimicrobial testing, bacteria were streaked to an LB agar plate and incubated for 24 h at 37 °C. Colonies from the resulting plates were used to start 3 ml LB broth cultures that were grown at 37 °C for 18 h overnight. To the ‘no peptide’ wells, 75 ul of sterile deionized water were added into the wells of tissue-culture treated 96-well polystyrene plates, and the remaining wells were set up as two-fold serial dilutions of each peptide starting at 256 ug/ml. Optical density of the bacterial cultures was measured at 600 nm (OD600) and cells were collected by centrifugation and normalized to an OD600 of 0.02 in LB and 75 ul of this suspension were aliquoted into the 75 ul of water or peptide solution. Thus, the final well contained 150 ul of liquid that was ½ strength LB and bacteria at a starting OD600 of 0.01. Identical plates were also generated with no bacteria added (only LB broth) to assess peptide aggregation to subtract from bacterial growth measurements. These 96-well plates were incubated with horizontal shaking at 120 rpm at 37 °C for 24 h and then OD600 was measured in a Biotek Synergy 2 plate reader.

The AMP cytotoxicity was tested with human A549 cells. The A549 cells was plated in 24-well plates at 300,000 cells/well. When cells were ~85% confluent, we added three different concentrations (30, 100, and 300 ug/ml) of the tested peptides for 24 hours. The cell viability was measured using a fluorescent dye calcein AM.

## Acknowledgments

We thank the Vermont Advanced Computing Core for supercomputing resources. J. L. and J. M. R were partially supported by an NIH R01 award (R01GM129431 to J. L.). J. B. F. were supported by an NSF grant (CHE-1945394 to J. L.). S.T.S. was supported by the U.S. Army Research Office (Grant 71015-CH-YIP).

