## Supplementary Info for "A Generative Approach toward Precision Antimicrobial Peptide Design"

**Table S1.** Comparison of sequence diversity, in terms of distribution of sequences of different E-values. The lower the E-value, the more likely the match is to be significant. E-values between 0.1 and 10 are generally dubious, and over 10 are unlikely to have biological significance.

| Generated peptides | 1 ≥ E-value | 10 ≥ E-value> 1 | E-value > 10 |
|---|---|---|---|
| AMP-GAN (8-10 residues) | 10% | 63% | 27% |
| CLaSS (< 20 residues, Ref. 1) | 52% | 30% | 18% |

**Table S2.** Generated sequences and their respective conditional labels. MIC50 label values were constant for all peptides and were in the lowest decile of the GAN effectively below 6.8 μg/mL.

| Sequence | Length (residue) | Target Microbe | Target Mechanism |
|---|---|---|---|
| SGRIASHFTQLWRWLRGYYKLM | 8 | Gram+, Gram- | Lipid Bilayer |
| QSGIFMHLKQLCRWLRGYMQWAGIG | 14 | Gram+, Gram- | Lipid Bilayer |
| QSNVFLSHFMQPCRWLRGKMG | 8 | Gram+, Gram- | Lipid Bilayer |
| MHKTQLRWFRCHLSQYPGAGL | 29 | Gram+, Gram-, Virus | Lipid Bilayer |
| WHLRRTRWRFEHLWSYGV | 14 | Gram+, Gram- | Lipid Bilayer |
| CSKWELRWRRYQGKVSYQLAL | 21 | Gram+ | Lipid Bilayer |
| QSRIFLSHFTQLSRWHRGKV | 12 | Gram+, Gram- | Lipid Bilayer |
| QSRIFLSHFYQPWRWLRGYMAWG | 17 | Gram+, Gram- | Lipid Bilayer |
| FHFTFLRWRRQHPYVQYQYGGIARAEML | 1 | Gram+, Gram- | Cytoplasmic Protein, Lipid Bilayer |
| QSRWLELAIQIRPNSGYQGIAGARLKRG | 8 | Gram+ | Lipid Bilayer |
| YHKMFLRWRFYQPYVSYQYGGIAGACGML | 9 | Gram+ | Lipid Bilayer |
| QSGIFPMHGKQLYRWHRGYMQWAG | 14 | Gram+, Gram- | Lipid Bilayer |

**Table S3.** Change in free energy (ΔG)* values for the simulated peptide sequences.

| Sequence | Free energy change (kcal/mol) |
|---|---|
| FTAAAEAEAAAA** | 27.0 (1.4)† |
| GIGKFLHSAKKFGKAFVGEIMNS*** | 20.1 (2.6)† |
| SGRIASHFTQLWRWLRGYYKLM | 18.5 |
| QSGIFMHLKQLCRWLRGYMQWAGIG | 21.8 |
| QSNVFLSHFMQPCRWLRGKMG | 23.7 |
| MHKTQLRWFRCHLSQYPGAGL | 24.6 |
| WHLRRTRWRFEHLWSYGV | 28.6 |
| CSKWELRWRRYQGKVSYQLAL | 28.6 |
| QSRIFLSHFTQLSRWHRGKV | 31.8 |
| QSRIFLSHFYQPWRWLRGYMAWG | 33.8 |
| FHFTFLRWRRQHPYVQYQYGGIARAEML | 35.6 |
| QSRWLELAIQIRPNSGYQGIAGARLKRG | 40.2 |
| YHKMFLRWRFYQPYVSYQYGGIAGACGML | 42.4 |
| QSGIFPMHGKQLYRWHRGYMQWAG | 47.0 |

* Calculated by the free energy of the umbrella window below the membrane minus the umbrella window above the membrane.
** The negative control sequence used to calculate $\Delta G_{-}$.
*** The positive control sequence used to calculate $\Delta G_{+}$.
† Errors (1s) for the control sequences are shown in parenthesis.

**Table S4.** Viability of cells as a percentage of control at different peptide concentrations.

| Pep1 | 30 µg/mL | 100 µg/mL | 300 µg/mL |
|---|---|---|---|
| Trial 1 | 100.0 ± 2.1 % | 105.7 ± 0.7 % | 4.3 ± 0.0 % |
| Trial 2 | 104.3 ± 3.8 % | 117.2 ± 2.3 % | 2.6 ± 0.0 % |
| Pep6 | 30 µg/mL | 100 µg/mL | 300 µg/mL |
| Trial 1 | 100 .0± 4.2 % | 95.6 ± 3.3 % | 95.5 ± 3.3 % |
| Trial 2 | 102.6 ± 2.6 % | 94.9 ± 1.5 % | 103.4 ± 3.1 % |

**Table S4.** Peptide helicity (%) measured from CD spectra. The secondary structure analysis was carried out with the program Dichromweb developed by Wallace et al. (1). Raw data of the CD spectra are provided in Figure S5.

| **% of helicity** | **Pep1** | **Pep2** | **Pep3** | **Pep4** | **Pep5** | **Pep6** |
|---|---|---|---|---|---|---|
| **w/o SDS** | 7 | 32 | 17 | 4 | 22 | 3 |
| **w SDS** | 73 | 80 | 58 | 15 | 6 | 5 |

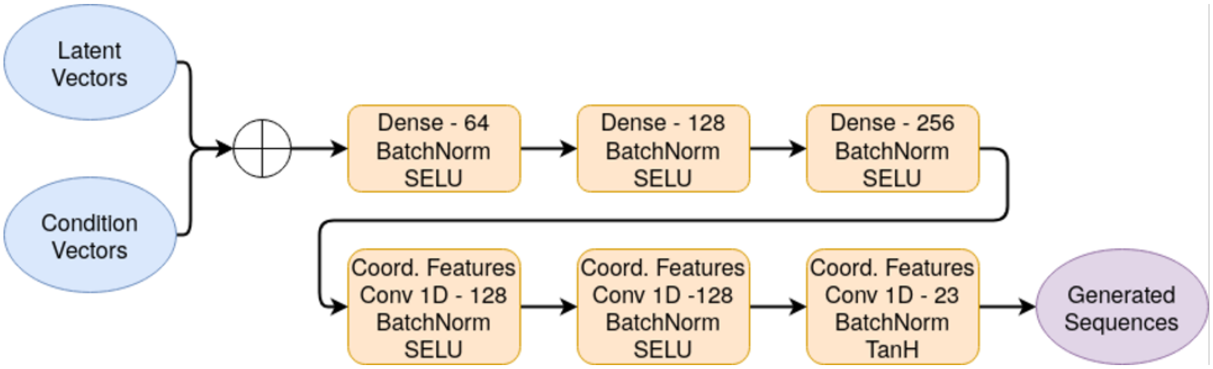

**Figure S1.** Details of the generator architecture used in AMP-GAN. Batch normalization is applied following each dense layer and convolution layer. The Scaled Exponential Linear activation function is applied to all layers, except the final layer that is terminated with a hyperbolic tangent activation.

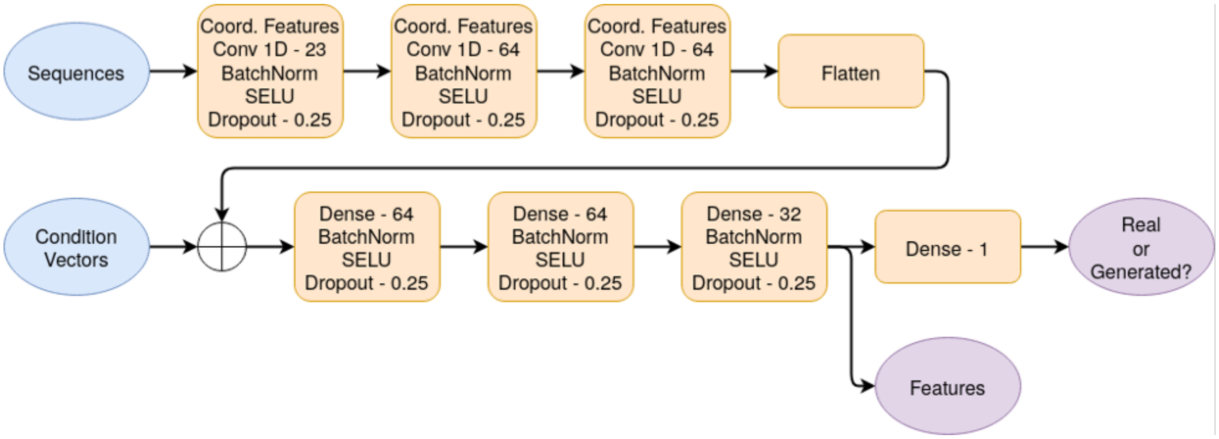

**Figure S2.** Details of the discriminator architecture used in AMP-GAN. Coordinate features are concatenated with the inputs to each convolution operation, providing global position information. Batch normalization is used to stabilize training and Dropout is applied to facilitate generator robustness.

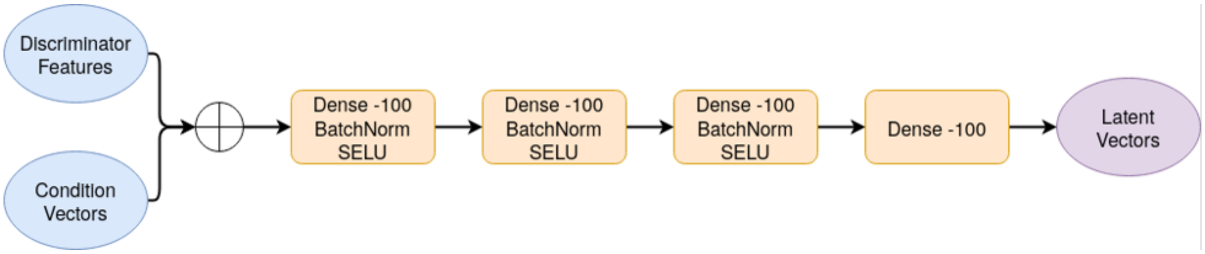

**Figure S3.** Details of the encoder architecture used in AMP-GAN.

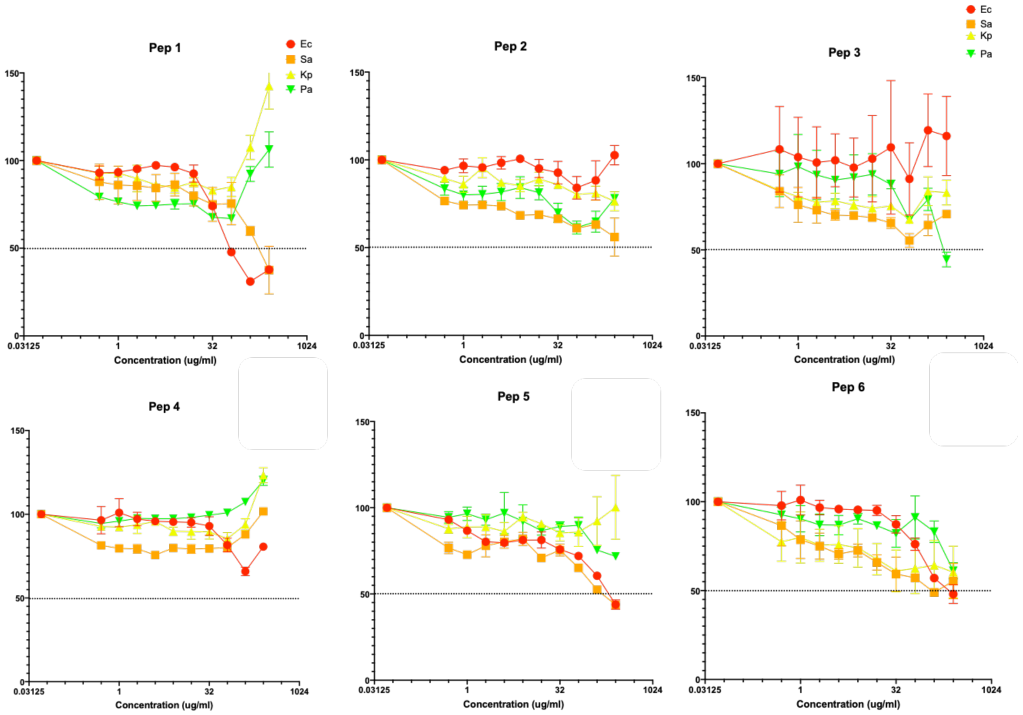

*Figure S4. Six novel peptides were tested to inhibit the growth of E. coli (red), S. aureus (orange), K. pneumoniae (yellow), and P. aeruginosa (green).*

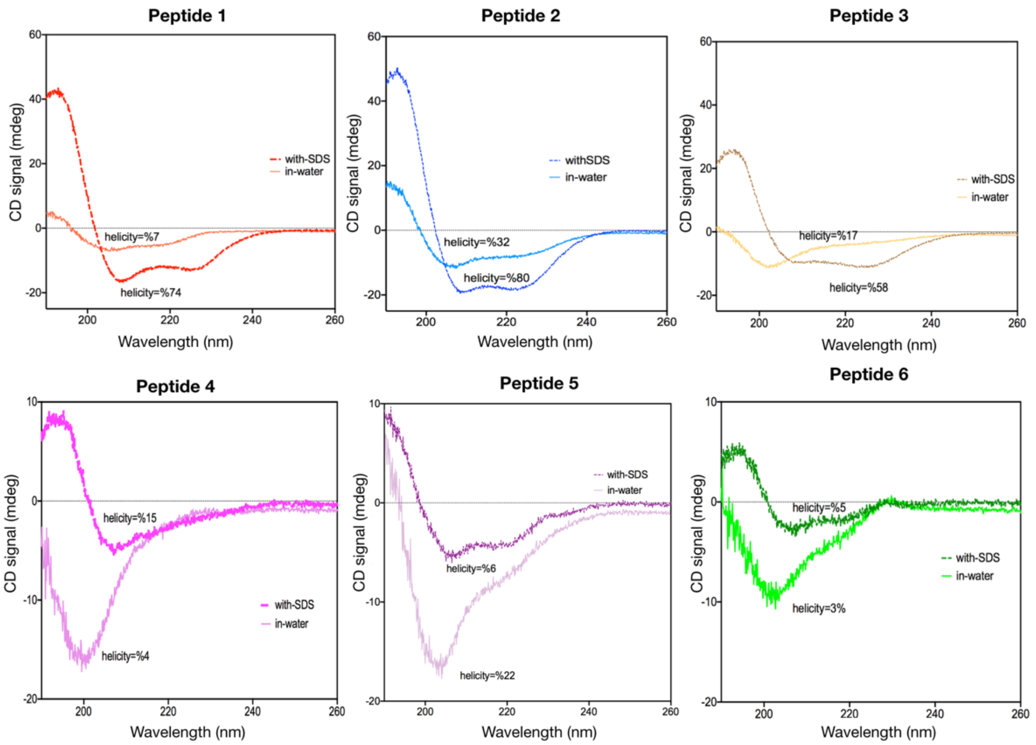

*Figure S5. CD spectra of all 6 peptides with and without SDS.*

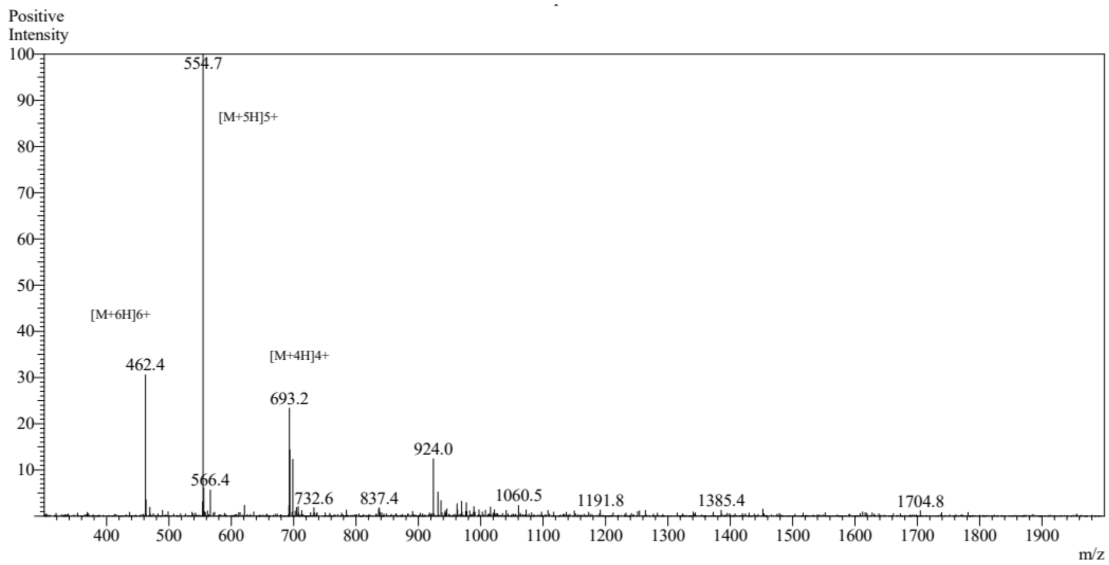

*Figure S6. Mass spectrum of Pep1.*

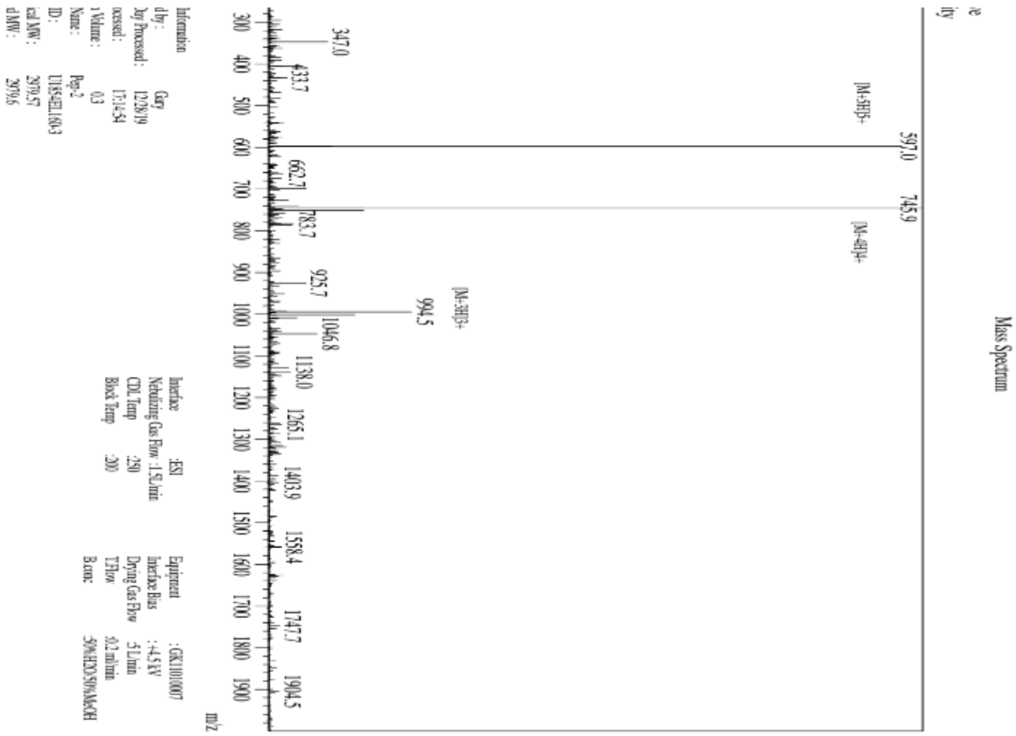

*Figure S7. Mass spectrum of Pep2.*

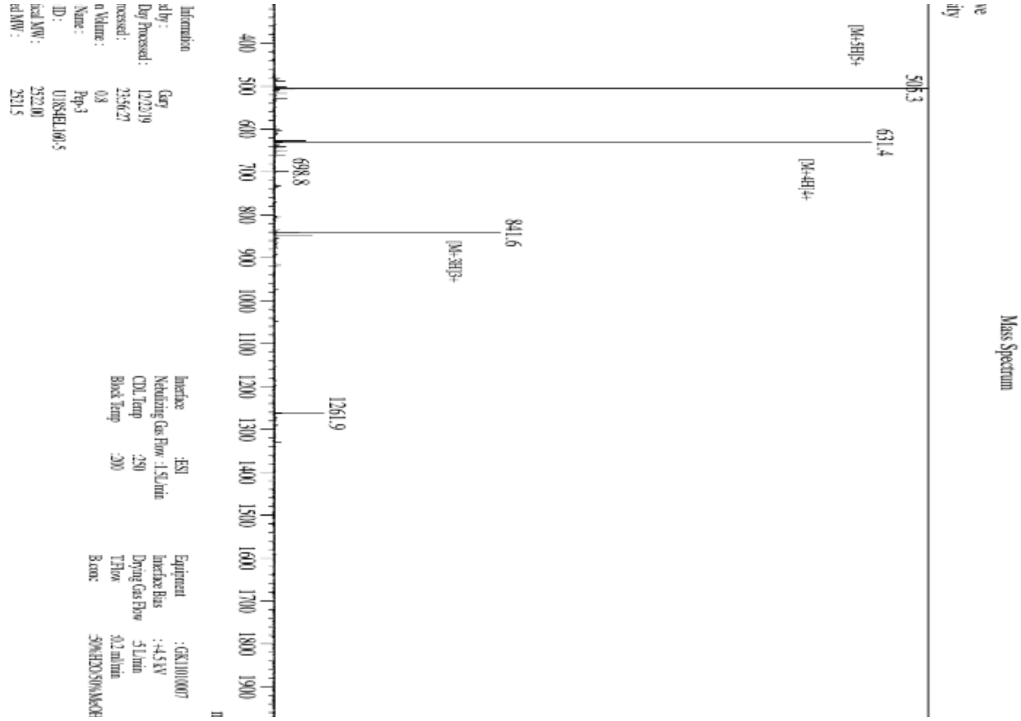

*Figure S8. Mass spectrum of Pep3.*

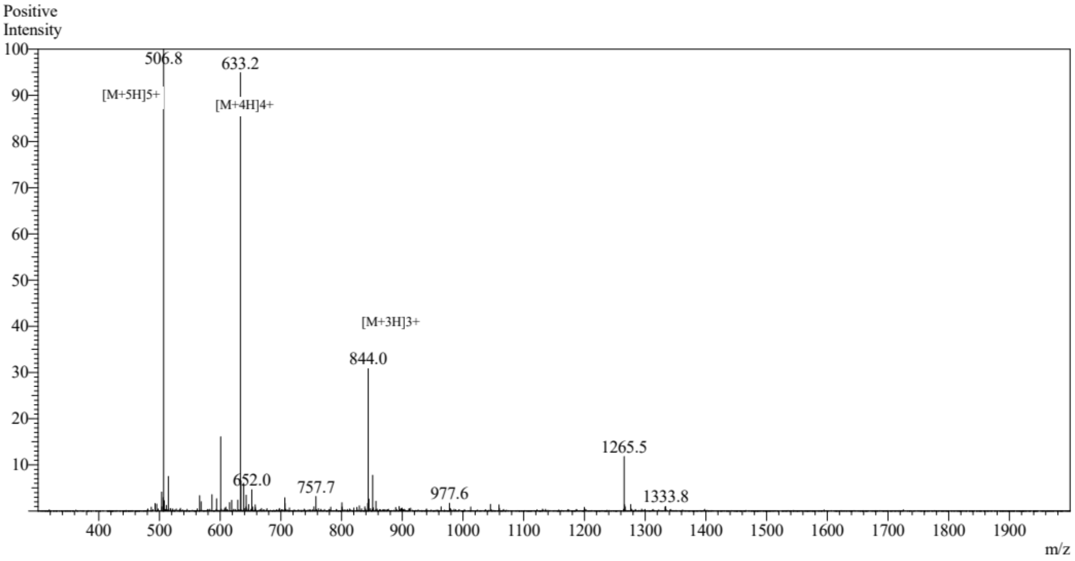

*Figure S9. Mass spectrum of Pep4.*

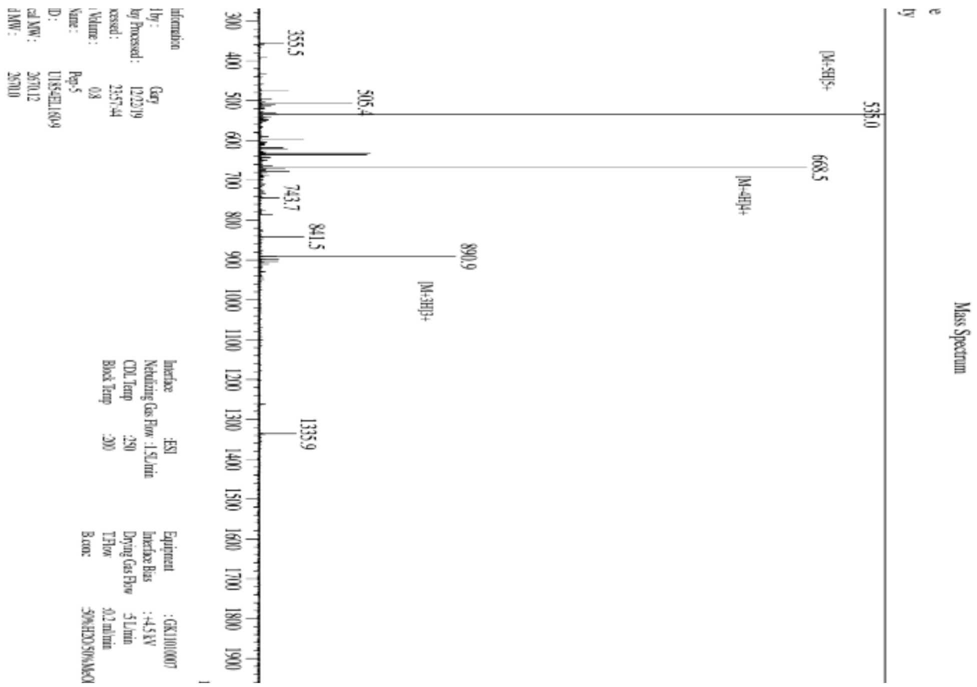

*Figure S10. Mass spectrum of Pep5.*

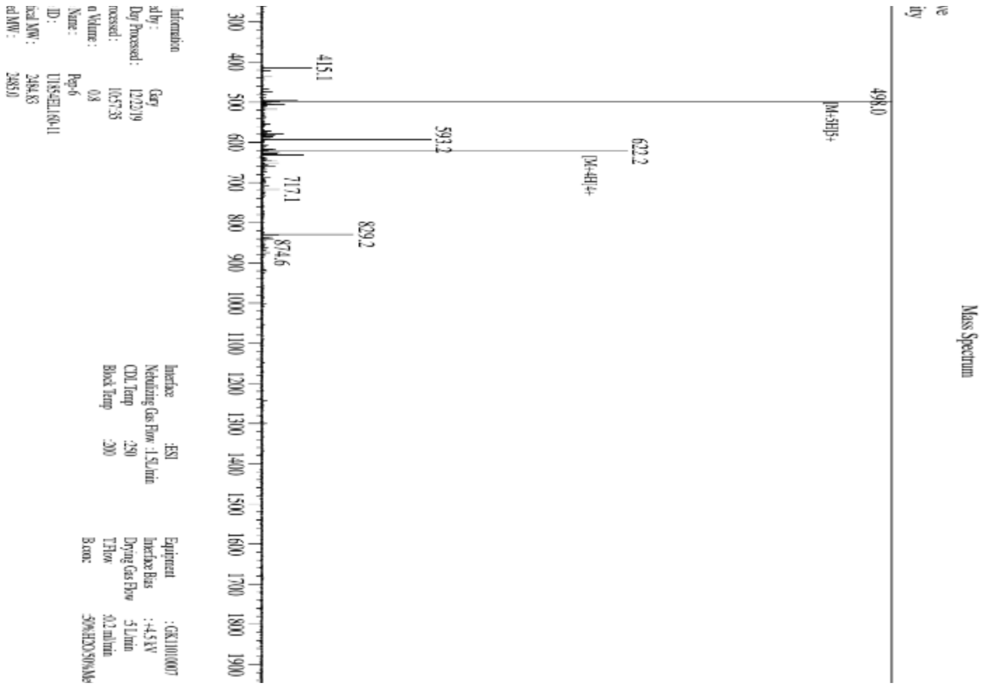

*Figure S11. Mass spectrum of Pep6.*
